# Acetyl transferase EP300 deficiency leads to chronic replication stress mediated by defective fork protection at stalled replication forks

**DOI:** 10.1101/2023.04.29.538781

**Authors:** Angelica Barreto-Galvez, Mrunmai Niljikar, Julia Gagliardi, Ranran Zhang, Vasudha Kumar, Aastha Juruwala, Archana Pradeep, Anam Shaikh, Priyanka Tiwari, Kritika Sharma, Jeannine Gerhardt, Jian Cao, Keisuke Kataoka, Adam Durbin, Jun Qi, B. Hilda Ye, Advaitha Madireddy

## Abstract

Mutations in the epigenetic regulator and global transcriptional activator, E1A binding protein (EP300), is being increasingly reported in aggressive hematological malignancies including adult T-cell leukemia/lymphoma (ATLL). However, the mechanistic contribution of EP300 dysregulation to cancer initiation and progression are currently unknown. Independent inhibition of EP300 in human cells results in the differential expression of genes involved in regulating the cell cycle, DNA replication and DNA damage response. Nevertheless, specific function played by EP300 in DNA replication initiation, progression and replication fork integrity has not been studied. Here, using ATLL cells as a model to study EP300 deficiency and an p300-selective PROTAC degrader, degrader as a pharmacologic tool, we reveal that EP300-mutated cells display prolonged cell cycle kinetics, due to pronounced dysregulations in DNA replication dynamics leading to persistent genomic instability. Aberrant DNA replication in EP300-mutated cells is characterized by elevated replication origin firing due to increased replisome pausing genome-wide. We demonstrate that EP300 deficiency results in nucleolytic degradation of nascently synthesized DNA at stalled forks due to a prominent defect in fork stabilization and protection. This in turn results in the accumulation of single stranded DNA gaps at collapsed replication forks, in EP300-deficient cells. Inhibition of Mre11 nuclease rescues the ssDNA accumulation indicating a dysregulation in downstream mechanisms that restrain nuclease activity at stalled forks. Importantly, we find that the absence of EP300 results in decreased expression of BRCA2 protein expression and a dependency on POLD3-mediated error-prone replication restart mechanisms. The overall S-phase abnormalities observed lead to under-replicated DNA in G2/M that instigates mitotic DNA synthesis. This in turn is associated with mitotic segregation defects characterized by elevated micronuclei formation, accumulation of cytosolic DNA and transmission of unrepaired inherited DNA lesions in the subsequent G1-phase in EP300-deficient cells. We demonstrate that the DNA replication dynamics of EP300-mutated cells ATLL cells recapitulate features of BRCA-deficient cancers. Altogether these results suggest that mutations in EP300 cause chronic DNA replication stress and defective replication fork restart results in persistent genomic instability that underlie aggressive chemo-resistant tumorigenesis in humans.

## Introduction

Loss-of-function mutations in EP300 and its close homolog, CBP, have been implicated in the pathogenesis of many hematologic malignancies^1^. EP300/CBP are global transcriptional coactivators that catalyze the addition of acetyl groups to lysine residues on histones (HAT activity) and non-histone proteins^2-5^ (KAT activity). Reports suggest that the independent inhibition of EP300 in human cells results in the differential expression of genes involved in regulating the cell cycle, DNA replication and DNA repair. While CBP downregulation results in changes to genes involved in antigen presentation, and terminal B-cell differentiation.

In this study, we sought to characterize the independent role of p300 in DNA replication integrity by studying patient-derived North American ATLL cell lines, that carry loss-of-function somatic mutations in *EP300* (but not *CBP*)^6^. In a recent retrospective analysis of a single center cohort of Caribbean/American ATLL (NA-ATLL), the median overall survival (OS) was only 6.9 months^7-10^, worse than the OS outcome in the largest ATLL cohort from Japan (J-ATLL) of ∼ 1 year^11^. Further analysis revealed that 20% of NA-ATLL patients had mutations in the EP300 gene, and that ATLL patients with epigenetic mutations had worse prognosis as compared to those without these mutations. Several patient-derived cell lines^12^ were subsequently developed that differ in EP300 mutation status. These cell line models provide a powerful tool to examine genotype to phenotype correlations associated with EP300 deficiency. To directly establish causal connections, we also took advantage of a highly selective EP300-specific PROTAC degrader that does not target CPB (a common problem with commercial EP300 inhibitors) and has limited toxicity in vivo^13^.

Importantly, almost all functional studies, in the literature, either examine p300 and CBP together or assess the importance of p300 to DNA repair since studies look at response to DNA damaging agents such as UV irradiation or ionizing radiation. For example, histone acetylation at double strand breaks (DSB) by p300/CBP has been shown to DNA repair proteins to chromatin^14-16^. In addition, numerous studies show that p300/CBP-mediated acetylation of proteins involved in DNA replication/repair can stimulate or inhibit their activities in reconstituted systems. These studies have revealed that p300/CBP can directly interact with PCNA **in vitro** to stimulate DNA synthesis after UV irradiation^17^, they can associate with ATR checkpoint signaling kinase^18^ and through their acetylation role (KAT), regulate the Fen1 endonuclease to help with Okazaki fragment maturation after UV irradiation^19^, DNA2 endonuclease^20^, DNA polymerase beta in base excision repair^21^, DNA glycosylases for base mismatch repair^22,23^, the Werner helicase after UV induced damage^24^ and numerous reports suggest their importance to homologous recombination repair^25^. Despite the suggested role for EP300 in DNA replication, it is not known whether inactivating mutations in EP300 spontaneously induce endogenous replication stress. Importantly, functional studies analyzing how these EP300Mut cells overcome replication fork collapse in the presence of replicative inhibitors such as aphidicolin and hydroxyurea is not known.

DNA replication integrity refers to the faithful and timely duplication of the genome to maintain genome stability. Replication stress (endogenous or induced), the biggest threat to genome stability, can stall the replisome and if not resolved efficiently, can lead to genomic instability. Upon stalling, human cells have evolved several mechanisms to stabilize and restart replication forks to prevent fork collapse leading to DNA breaks. Cells activate the ATR/Chk1 checkpoint kinase signaling pathway, in response to replication stress. Phosphorylation of ATR, Chk1, RPA, and histone H2AX then ensure that DNA replication stalls in order to resolve stalled replication forks before replication can continue. Fork restart is then mediated by a complex set of steps that involve fork reversal by SMARCAL1, HTLF, ZRANB3 and FBH1, fork stabilization by the RAD51 recombinase downstream mediators such as BRCA2/BRCA1/PALB2 and fork restoration by error-free recombination-based or error-prone POLD3 mediated-mechanisms. While p300 has been implicated in transcriptionally activating BRCA1 and RAD51 to promote homologous recombination^25,26^, the importance of p300 acetylation and possible transcriptional control for the downstream mediators of replication fork stability in response to replication stress has not been evaluated.

To address p300’s role in DNA replication, we analyzed replication integrity, cell cycle progression and evaluated mechanisms driving genomic instability in p300-defeicient cells. We found that EP300-mutated cells display a protracted cell cycle, extensive genomic instability, dysregulations in DNA replication and repair dynamics. Using a powerful locus specific single molecule assay and genome-wide DNA fiber analysis, we show that perturbed DNA replication in EP300-mutated ATLL cells is characterized by highly elevated replication origin firing, increased replisome pausing and prominent nucleolytic degradation of nascent DNA at collapsed forks. This in turn results in the persistence of single stranded DNA (ssDNA) genome-wide, resulting in the hyperphosphorylation of RPA at Ser4/8, a prominent parker of collapsed replication forks, in EP300-deficient NA-ATLL cells. Inhibition of Mre11 nuclease rescued the ssDNA accumulation indicating a dysregulation in downstream mechanisms that restrain nuclease activity at stalled forks. To address isogeneity, we have utilized a highly selective EP300-specific PROTAC degrader, JQAD1, that does not target CPB (a common problem with commercial EP300 inhibitors) and has limited toxicity in vivo^13^. These analyses recapitulate phenotypes observed in EP300Mut patient cells. Taken together, these results reveal that p300 deficiency closely resembles BRCA-deficient cancers. Elevated endogenous genomic instability, observed in ATLL cells during the S-phase is exacerbated by EP300 deficiency, even in the absence of exogenous stress/damage. These S-phase abnormalities result in under-replicated DNA in G2/M, instigate mitotic DNA synthesis, and are associated with mitotic segregation defects, elevated micronuclei, and accumulation of cytosolic DNA in NA-ATLL cells.

## Results

### *EP300*-mutated cells have decreased histone acetylation, aberrant cell cycle dynamics, spontaneous DNA damage and robust replicative checkpoint activation

Acetylation on histone H3 lysine 27 (H3K27ac), a marker of master transcriptional regulation loci^27,28^, is catalyzed by p300 and its paralogue CBP^29^. To study H3K27 acetylation in EP300Mut cells, we carried out immunoblotting analysis of protein extracts from ATLL cells lines differing in EP300 mutational status. Absence of EP300 corresponds to a significant decrease in the H3K27ac mark genome-wide in ATLL cells with mutated EP300 status (**Figure 1A**). These results are further confirmed by the decrease in the H3K27ac mark observed in EP300WT cells exposed to the PROTAC-mediated EP300-specific degrader JQAD1 (**Figure 1B)**.

**Figure 1:**
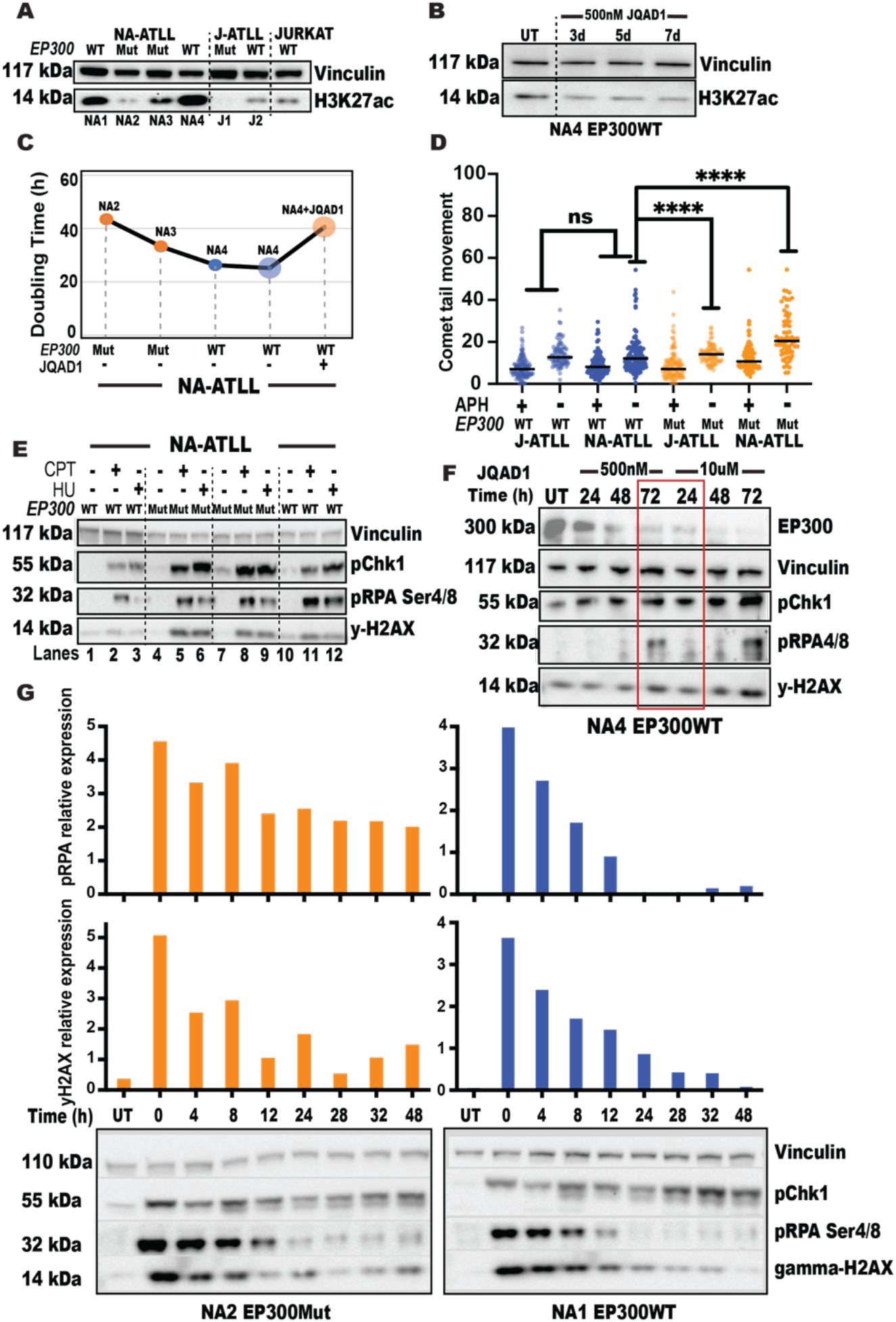
*EP300*-mutated cells have decreased histone acetylation, aberrant cell cycle dynamics, spontaneous DNA damage and robust replicative checkpoint activation. A. Expression levels of H3K27 histone acetylation mark by western blotting in EP300WT and EP300Mut NA/J-ATLL cells and Jurkat cells. Expression levels of Vinculin was used as a loading control; B. Expression levels of H3K27 histone acetylation mark by western blotting after p300 protein degradation using JQAD1 PROTAC compound; C. Calculation of cell doubling times using the open source cell doubling time calculator (https://www.omnicalculator.com/biology/cell-doubling-time). Cell doubling was calculated every 48 hours from three independent cultures of cells; D. Measurement of DNA single strand breaks by alkaline Comet assay in EP300Mut and EP300WT cells treated with Aphidicolin (APH). Comet tail lengths were measured using the OpenComet plugin as part of the ImageJ software, n=150; E-F. Expression levels of p300 protein, phospho-Chk1 Ser317, phospho-RPA Ser4/8 and phospho-Histone H2AX Ser139 by western blotting. Expression levels of Vinculin was used as a loading control; and G. Time course experiment to measure recovery of cells after release into drug-free media over the course of 48hours. Cells were collected at eight time points (0, 4, 8, 12, 24, 28, 32 and 48hours) and expression levels of phospho-Chk1 Ser317, phospho-RPA Ser4/8 and phospho-Histone H2AX Ser139 were measured by western blotting. Expression levels of Vinculin was used as a loading control. The p-values are indicated as follows: * <0.03, ** <0.0021, *** <0.0002, **** <0.0001. Scale bar 10 μm.

To assess the effect of EP300 deficiency on cell cycle progression, we analyzed cell doubling time in EP300Mut, EP300WT and EP300WT+JQAD1 cell lines without exogenous replication stress. These results revealed that in the absence of EP300, cells take longer to progress though cell cycle, as compared to EP300WT cells (**Figure 1C; Supplemental Figure 1A**). One of the hallmarks of cancer is genomic instability resulting from spontaneous DNA replication stress^30^. To understand whether there is an EP300 dependent increase in genomic instability in NA-ATLL cells, we measured the overall magnitude of spontaneous and induced genomic instability by immunostaining (IF) for phosphorylation of histone variant 2A (γH2AX). This analysis revealed a spontaneous significant increase in γH2AX foci per cell in NA-ATLL EP300Mut cells and this effect was further exacerbated upon replicative inhibition (**Supplementary Figure 1B-C**). We then validated instability at the single cell level by measuring DNA breaks using the Comet assay^31^. Similar to the IF results, we observed a significant increase in comet tail movement in EP300Mut NA-ATLL cells (orange), compared to the EP300WT cells (Blue-**Figure 1D; Supplementary Figure 1D**).

Since replicative difficulties during the S-phase results in activation of ATR-Chk1 signaling cascade^32^, we next evaluated checkpoint activation by immunoblotting in EP300Mut and WT cells treated with hydroxyurea (HU, replicative inhibitor) or camptothecin (CPT, Topoisomerase 1 inhibitor). While inhibitor treatments induced robust Chk1 phosphorylation in all cell lines, EP300 deficiency (EP300Mut) was associated with stronger pChk1 signal (**Figure 1E-F**), even in the absence of replication inhibitors, indicating increased endogenous replication stress in the absence of p300. Chk1 phosphorylation was further exacerbated upon replicative inhibition. To establish that replication arrest mediated by Chk1 phosphorylation is in response to replication perturbation, we assessed phosphorylation of RPA at Ser 4/8 (pRPA4/8), a marker of collapsed forks and ssDNA^33^, that is critical for RAD51 recruitment and fork restart following HU^33^. Again, while all cell lines activated pRPA4/8 in response to damage, EP300Mut cells hyperphosphorylated RPA, indicating an increased number of collapsed forks and ssDNA in the absence of p300 protein.

### EP300-mutated NA-ATLL cells have persistent checkpoint signaling associated with unresolved DNA damage

While checkpoint activation appears to be proficient and even robust in EP300Mut cells, the slow s-phase progression indicated by longer doubling times and extensive DNA damage point to a possible defect in damage and checkpoint resolution before G2/M transition. To assess this, we carried out a time course experiment in EP300Mut and WT NA-ATLL lines. The results from this analysis showed a clear separation in damage and checkpoint resolution capabilities between the EP300Mut and EP300WT cohorts (**Figure 1G; Supplemental Figure 1E)**. While phosphorylation of histone variant 2A (γH2AX), a marker of DNA damage and pRPA4/8 are efficiently resolved in the EP300WT NA-ATLL cells by 24 hours (blue bars), EP300Mut cells (orange bars) never completely resolve the damage. These studies show that EP300 deficiency leads to aberrant cell cycle progression due to spontaneous endogenous stress, increased sensitivity to exogenous replication inhibition and persistent DNA damage and replicative checkpoint activation.

### EP300 mutated cells have aberrant replication dynamics associated with increased origin firing

To understand the endogenous events driving spontaneous replication stress in EP300Mut ATLL cells, we next investigated the DNA replication dynamics. Chromosomal instability in NA-ATLLL cells has been shown to cluster at difficult to replicate regions of our genome called common fragile sites (CFS)^34-36^, identifying CFS as endogenous hotspots of replicative distress. Studying DNA replication dynamics at a specific genomic locus would eliminate the need for exogenous inhibitor treatments and enable us to understand the specific effect of protein deficiencies, in this case p300, and the associated changes to the cellular microenvironment on DNA replication. We are uniquely equipped to study locus specific DNA replication using our powerful high-resolution

DNA replication assay, **S**ingle **M**olecule **A**nalysis of **R**eplicated **D**NA (SMARD; **Supplemental Figure 2A**). To understand the effect of EP300 deficiency on in vivo DNA replication dynamics, we used SMARD analysis to establish replication programs at common fragile site FRA6E (CFS-FRA6E; **Figure 2A**), in pairs of NA-ATLL and J-ATLL cells differing in EP300 mutational status. Under normal conditions, the replication program at CFS-FRA6E in an EP300WT cell line is predominantly replicated by forks travelling in the 5’ to 3’ direction and there are no major replication origins within the locus^37^. The absence of replication origins is a defining characteristic of CFS loci and we have previously established that any origins observed is a stress response either due to the absence of an essential protein^38^ or due to exposure to replicative inhibitors^39^. Replication dynamics at FRA6E in a J-ATLL or NA-ATLL cell line expressing wildtype EP300 revealed a replication program very similar to the untreated lymphocyte FRA6E profiles previously characterized by our lab^37^. The FRA6E locus was replicated by forks primarily moving in the 5’ to 3’ direction and while there were no prominent replication initiation sites to be detected with the 375 kilobase region analyzed, EP300WT NA-ATLL cells showed some replication pausing, likely due to some endogenous stress (**Figure 2B,D**). However, EP300Mut ATLL cells revealed significant replication perturbation characterized by a striking increase in dormant origin activation (**Figure 2C,E**-red box**; Figure 2F**-orange bars), replication pausing (white rectangles) and replication fork directionality change (**Figure 2C,E**). These results suggest that lack of p300 expression results in spontaneous replication stress that dysregulates DNA replication dynamics at the most vulnerable regions of our genome, fragile sites.

**Figure 2:**
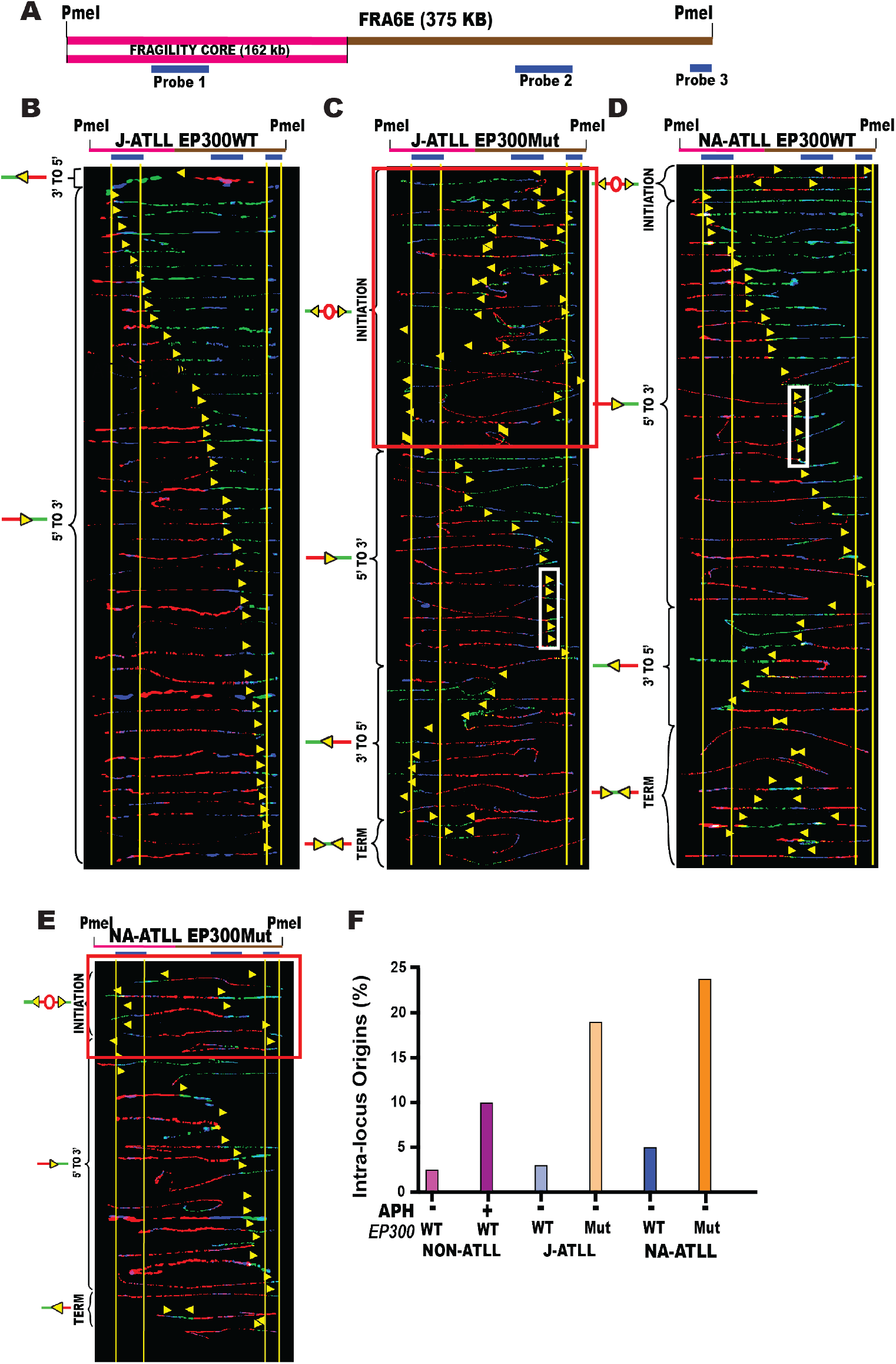
Aberrant replication dynamics in EP300Mut ATLL cells. (A) Locus map of a 375 kb region in the CFS-FRA6E obtained by *PmeI* digestion. The region includes the fragility core of CFS-FRA6E (pink line – 162 kb). The FISH probes that identify the segment are labeled in blue. **Top**; Locus map of the *PmeI* digested FRA6E segment. **Bottom**; Aligned photomicrograph images of labeled DNA molecules from DNA Replication program at CFS-FRA6E in (B) J-ATLL EP300WT (J2); (C) J-ATLL EP300Mut cells (J1); (D) NA-ATLL EP300WT cells (NA1); and (E); NA-ATLL EP300Mut (NA2) cells. The yellow arrows indicate the sites along the molecules where the IdU transitioned to CldU. The molecules are arranged in the following order: molecules with initiation events, molecules with 3’ to 5’ travelling forks, molecules with 5’ to 3’ travelling forks and molecules with termination events; (F) Percentage of molecules with initiation sites in FRA6E in J-ATLL EP300WT, J-ATLL EP300Mut, NA-ATLL EP300WT and NA-ATLL EP300Mut cells.

### EP300 deficiency results in genome-wide replication stalling and fork collapse

To understand whether spontaneous replication stress observed in the absence of p300 impacts replication dynamics genome-wide, we measured replication fork stalling in EP300Mut and WT cells using DNA fiber analysis^40^. Here, cells were pulse labelled with nucleoside analog IdU for 30 mins following by CldU for 60 mins during which a replicative inhibitor is added to assess fork stalling. A shortening of the CldU tract (indicated by a ratio <3) would be indicative of replication stalling. Results from this analysis showed that HU treatment resulted in replication fork stalling in all the cell lines analyzed. However, a comparison of the extent of stalling between the treated pairs revealed significantly shorter CldU lengths in EP300Mut cell lines (orange dots) as compared to EP300WT lines (**Figure 3B**; blue dots). Significant increased replication stalling due to EP300 deficiency was confirmed in a JQAD1 treated EP300WT cell line. In contrast, EP300WT lines from each cohort did not differ significantly in CldU tract lengths, further highlighting the EP300 specificity of this phenotype (**Figure 3B**). Cells employ specific mechanisms to overcome fork stalling/pausing to ultimately prevent fork collapse, where the replication fork loses its ability to synthesize DNA^41^. Upon fork stalling, cells stabilize the stalled fork by a prominent mechanism called fork regression or reversal (**Figure 3A**)^42,43^. Failure to repair and restart the reversed forks results in nucleolytic degradation of nascent regressed DNA^44^. To assess nucleolytic degradation, cells are first pulse labelled with IdU for 30 mins followed by CldU for 30 mins and then treated with HU for 4 hours in the absence of analogs. A shortening of the CldU tract would be indicative of fork degradation. Analysis of NA-ATLL cell lines differing in EP300 mutational status showed that there was no significant nascent strand degradation at stalled forks in EP300WT cells (**Figure 3C**; blue dots). However, EP300Mut cells displayed a significant increase in nucleolytic degradation, as compared to untreated cells (**Figure 3C**; orange dots), likely indicating a prominent fork restart defect in the absence of EP300.

**Figure 3:**
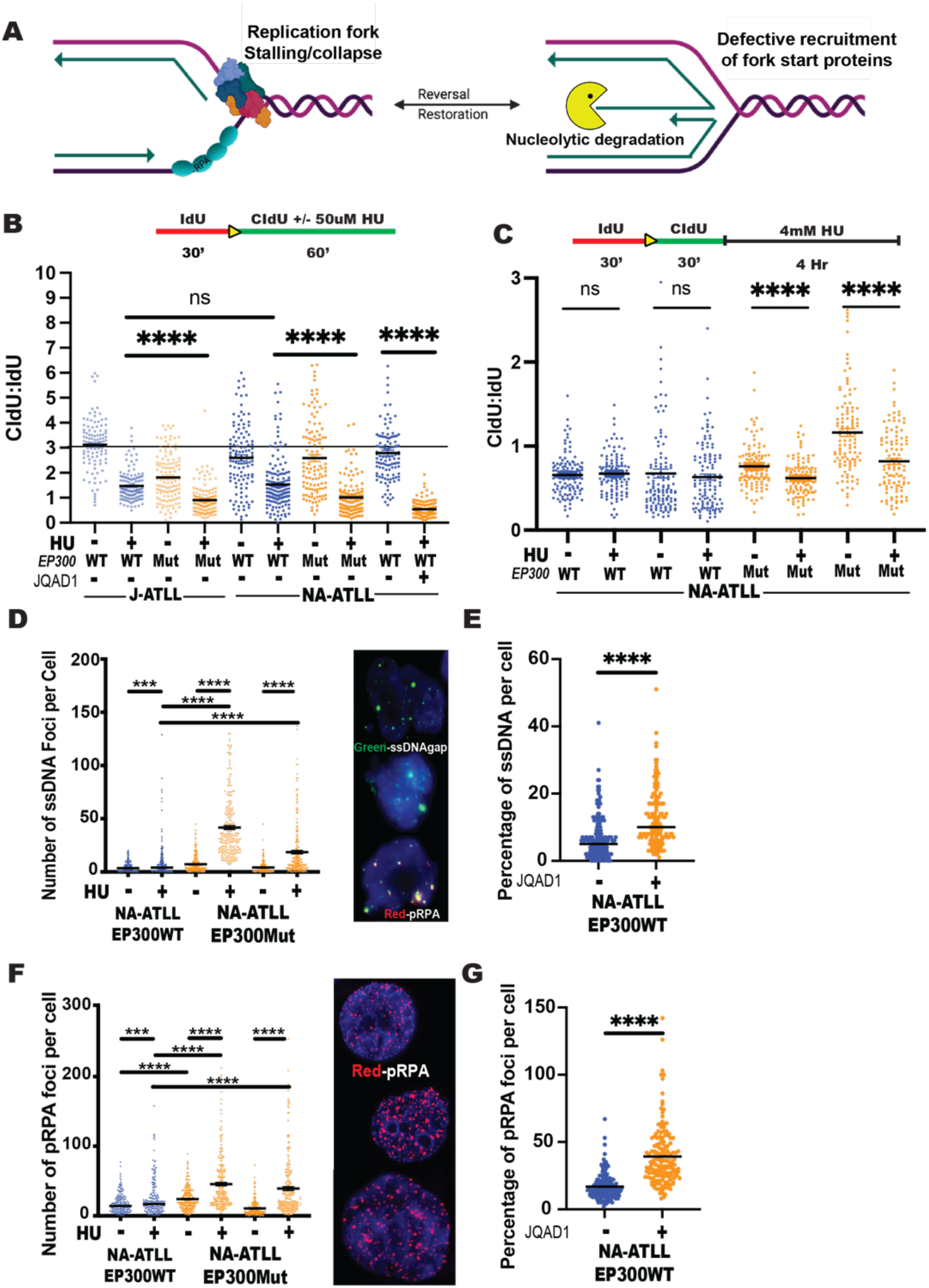
EP300 deficiency results in genome-wide replication stalling and fork collapse, leading to accumulation of extensive ssDNA gaps. A. Schematic model depicting replication fork stalling, fork protection and nucleolytic degradation; B. DNA fiber analysis of 50uM hydroxyurea (HU) treated EP300WT (J2, NA1), EP300Mut (J1, NA2) and EP300WT+JQAD1 treated NA-ATLL (NA4) cells to assess replication fork stalling. The fork rate (CldU/IdU ratio) is indicated, n=150. The p-values are indicated as follows: * <0.03, ** <0.0021, *** <0.0002, **** <0.0001. Scale bar 10 μm; C. DNA fiber analysis measuring nucleolytic degradation after 4mM HU treatment in EP300WT, EP300Mut NA-ATLL cells. The fork rate (CldU/IdU ratio) is indicated, n=150. The p-values are indicated as follows: * <0.03, ** <0.0021, *** <0.0002, **** <0.0001. Scale bar 10 μm; D-E. Analysis of number of single stranded gaps/breaks (ssDNA)/ Iododeoxyuridine (IdU) foci (green) per cell nuclei (DAPI, blue) in EP300Wt (NA4) and EP300Mut (NA2, NA3) cells exposed to HU. ssDNA gaps are measured by IdU incorporation under non-denaturing conditions, representative images are shown on the right, n=150; E. Analysis of number of ssDNA/ IdU foci (green) per cell nuclei (DAPI, blue) in EP300WT cells treated with JQAD1 PROTAC compound that degrades p300 protein; F. Analysis of number of pRPA Ser4/8 foci (red) per cell nuclei (DAPI, blue) in EP300Wt (NA4) and EP300Mut (NA2, NA3) cells exposed to HU, representative images are shown on the right, n=150; G. Analysis of number of pRPA Ser4/8 foci (red) per cell nuclei (DAPI, blue) in in EP300WT cells treated with JQAD1. The p-values are indicated as follows: * <0.03, ** <0.0021, *** <0.0002, **** <0.0001. Scale bar 10 μm.

### Fork restart defects in the absence of EP300 result in the accumulation of extensive ssDNA gaps

Fork stalling and nucleolytic degradation of nascent strands can lead to extensive ssDNA breaks/gaps in the genome. To assess ssDNA accumulation in the absence of EP300, we assessed BrdU incorporation under non-denaturing conditions by immunofluorescence staining (green foci in representative nuclei). The results showed a striking increase in the number of ssDNA foci detected per cell in EP300Mut NA-ATLL cells (**Figure 3D**; orange dots). This effect that was significantly higher in EP300Mut cells even when compared to treated EP300WT cells (blue dots). Inhibitor treatment in EP300Mut cells exacerbated the ssDNA accumulation even further. Importantly, JQAD1-mediated p300 degradation in EP300WT cells resulted in a significant increase in spontaneous ssDNA gaps, clearly establishing that ssDNA gaps were being generated due to the absence of p300 (**Figure 3E**; JQAD1-orange dots). To establish that the source of the ssDNA were indeed stalled/collapsed forks, we co-stained for pRPA Ser4/8. Similar to the ssDNA data, these results revealed a striking increase the number of pRPA4/8 foci detected per cell in EP300Mut NA-ATLL cells (orange dots), compared to EP300WT cell line (**Figure 3F**; blue dots). Similar results were obtained upon p300 degradation in a EP300WT cell line (**Figure 3G**; JQAD1-orange dots). These results collectively establish collapsed forks as the primary sites of ssDNA gap/break formation in the absence of p300 protein expression.

The most common nucleases implicated in nucleolytic processing of nascent DNA are Mre11, DNA2 and Exo1^45^. To determine the mechanistic basis for extensive ssDNA gap/break, marked by hyperphosphorylated RPA at Ser4/8, in EP300Mut cells, we treated cells with Mirin, a potent inhibitor of Mre11 activity in the presence or absence of HU. The immunoblotting analysis of whole cell lysates showed that the hyperphosphorylation of RPA observed largely in the EP300Mut NA-ATLL cells is almost completely rescued by Mre11 inhibition (**Figure 4A**; orange bars). Mirin treatment of EP300WT N-ATLL cells did not result in any significant change (**Figure 4A**; blue bars). Importantly, immunofluorescence staining for BrdU under non-denaturing conditions after Mirin treatment resulted in a highly significant reduction in ssDNA gap/break formation in EP300Mut cells (orange dots) as compared to EP300WT cells (**Figure 4B**; blue dots). These results clearly establish that the formation of ssDNA gap/breaks in the absence of EP300 is driven by the hyperactivity of Mre11 nuclease likely due to a prominent defect in downstream fork restart machinery.

**Figure 4:**
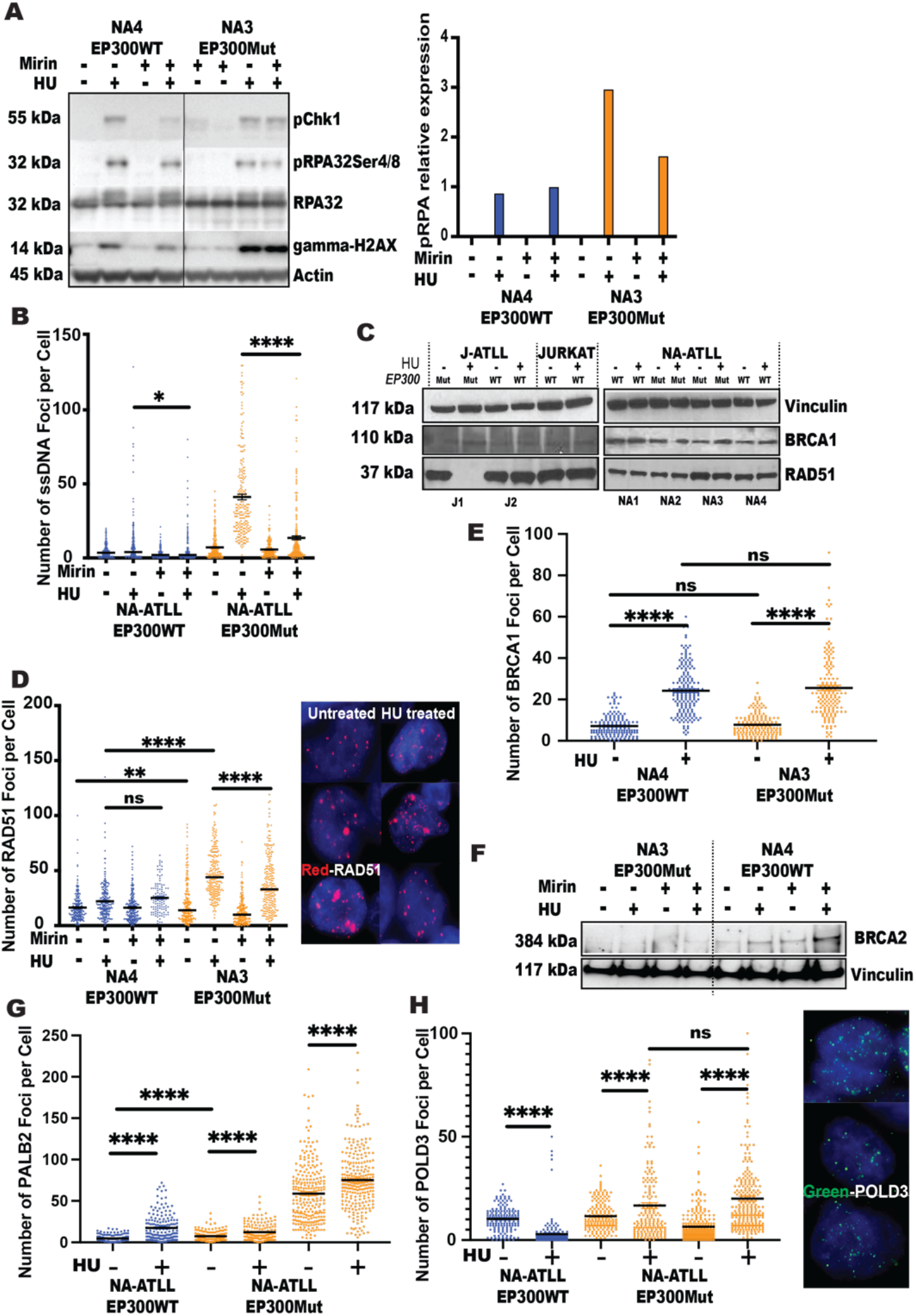
EP300 deficient cells have a prominent defect in downstream fork restart machinery. A. Expression levels of phospho-Chk1 Ser317, phospho-RPA Ser4/8 and phospho-Histone H2AX Ser139 in EP300WT (NA1) and EP300Mut NA-ATLL (NA2) cells treated with HU, in the presence of absence of Mirin, by western blotting. Expression levels of Actin was used as a loading control; B. Analysis of number of single stranded gaps/breaks (ssDNA)/ Iododeoxyuridine (IdU) foci per cell nucleus (DAPI, blue) in EP300Wt (NA1) and EP300Mut (NA2) cells exposed to HU, in the presence or absence of Mirin (Mre11 nuclease inhibitor). C. Expression levels of BRCA1 and RAD51 proteins from WCE in EP300WT and EP300Mut J/NA-ATLL cells and Jurat cells treated with HU, by western blotting. Expression levels of Vinculin was used as a loading control; D. Analysis of number of RAD51 foci (red) per cell nucleus (DAPI, blue) in EP300Wt and EP300Mut cells exposed to HU, in the presence or absence of Mirin, representative images are shown on the right, n=250; E. Analysis of number of BRCA1 foci per cell nucleus (DAPI, blue) in EP300Wt and EP300Mut NA-ATLL cells exposed to HU, n=250; F. Expression levels of BRCA2 protein from WCE in EP300WT and EP300Mut NA-ATLL cells treated with HU, in the presence or absence of Mirin, by western blotting. Expression levels of Vinculin was used as a loading control; G. Analysis of number of PALB2 foci per cell nucleus (DAPI, blue) in EP300Wt and EP300Mut NA-ATLL cells exposed to HU, n=250; H. Analysis of number of POLD3 foci (green) per cell nucleus (DAPI, blue) in EP300Wt and EP300Mut NA-ATLL cells exposed to HU, representative images are shown in the right, n=150. The p-values are indicated as follows: * <0.03, ** <0.0021, *** <0.0002, **** <0.0001. Scale bar 10 μm.

### EP300 deficient cells have a prominent defect in downstream fork restart machinery

Extensive nucleolytic processing of stalled forks by nucleases has been previously described in the absence of downstream fork restart machinery (BRCA1-BRCA2-PALB2-RAD51)^44^. Partial inhibition of RAD51 has been shown to induce nascent strand degradation in response to DNA damaging agents^46,47^. Importantly, p300 has been implicated in regulating BRCA1 and RAD51 expression to promote homologous recombination^25,26^. To address this, we first carried out immunoblotting analysis to measure BRCA1 and RAD51 protein expression from whole cell extracts (WCE) in EP300Mut and EP300WT cells. The results showed no differences in RAD51 or BRCA1 protein expression levels irrespective of EP300 mutational status (**Figure 4C**). In addition, independent degradation of p300 protein in EP300WT cells by JQAD1 did not alter BRCA1 expression in cells (**Supplementary Figure 4A**). Next, we assessed chromatin recruitment and loading of RAD51 nucleofilaments by immunofluorescence staining. The results showed that while RAD51 recruit to chromatin and loading were present in both EP300Mut and EP300WT cells, EP300Mut cells displayed increase spontaneous RAD51 chromatin loading, consistent with the spontaneous increase in ssDNA observed in these cells (**Figure 4D;** red foci in representative images). In addition, RAD51 recruitment was significantly increased after HU treatment, especially in the absence of p300. Interestingly, while Mre11 degradation partially reduced RAD51 loading in EP300Mut cells (orange dots +Mirin +HU), no difference was observed in EP300WT cells (**Figure 4D;** blue dots +Mirin +HU). These results clearly indicate that elevated RAD51 recruitment in EP300Mut cells, beyond what is observed in EP300WT cells is due to increased Mre11 generated ssDNA at collapsed replication forks. The Mre11 activity and efficient RAD51 loading also likely indicate that fork regression is quite proficient in the absence of p300.

Nucleolytic degradation has also been described in the absence of downstream effectors of fork protection, such as BRCA1, BRCA2, PALB2 and FANCD2^44^. While BRCA1 protein expression is proficient in EP300Mut cells, we carried out an immunofluorescence analysis to measure BRCA1 chromatin recruitment. The results revealed that while HU treatment increased BRCA1 foci formation in general, similar levels of BRCA1 foci per cell were detected in both EP300WT and EP300Mut cells (**Figure 4E**). Fork degradation has been extensively associated with BRCA2 deficiency^48,49^. To assess BRCA2 expression, we carried out immunoblotting of WCE from EP300WT and EP300Mut cells. The results revealed a prominent downregulation in BRCA2 expression in the absence of p300 expression (**Figure 4F**). In contrast, BRCA2 expression was present in EP300WT cells, increased upon HU treatment and wasn’t significantly altered by Mre11 inhibition. These results collectively indicate that a downregulation of BRCA2 protein expression in EP300 deficient cells might be the underlying mechanism for defective replication fork stability and integrity. The mechanisms underlying BRCA2 downregulation need further assessment and evaluation.

### Error-prone repair mechanisms are upregulated in EP300 mutated NA-ATLL cells

It has been reported that breakage at replication forks in stressed cells that are deficient in proteins involved in fork protection and HR are repaired by Microhomology mediated Break Induced Replication (MMBIR) repair pathway^50,51^. The MMBIR pathway requires POLD3, and Rad52 instead of PALB2, BRCA1/2 and RAD51. To assess dependency on error-prone versus error-free replication restart in EP300Mut cells, we analyzed recruitment of PALB2, as essential factor for fork restart and HR and POLD3, which is an essential factor of the MMBIR repair pathway^52,53^ by immunofluorescence staining. Assessment of spontaneous PALB2 foci formation revealed a significant increase in EP300Mut NA-ATLL cells as compared to NA-ATLL EP300WT cells (**Figure 4G; Supplementary Figure 4F**). Upon HU treatment, all cell lines showed an increase in PALB2 foci formation irrespective of p300 expression (**Figure 4G; Supplementary Figure 4B**). Interestingly, treatment with an alternate replicative inhibitor aphidicolin (DNA polymerase inhibitor) in low doses, resulted in a significant decrease in PALB2 chromatin recruitment in EP300Mut ATLL cells, compared to their untreated pairs (**Supplementary Figure 4E-F**). Analysis of POLD3 chromatin recruitment to assess dependency of EP300Mut ATLL cells on error-prone restart after fork stalling revealed a significant increase in POLD3 foci per cell in EP300Mut J-ATLL and NA-ATLL cell lines, as compared to EP300WT cells (**Figure 4H; Supplementary Figure 4C**). Importantly, Mre11 inhibition led to a significant decrease in the POLD3 foci per cell EP300Mut NA-ATLL cells (**Supplementary Figure 4D**), showing the error-prone repair was the prevalent mechanism of replication-associated repair of collapsed forks in these cells. These results provide early evidence of upregulation of error-prone fork restart and repair, such as the MMBIR mechanisms, in EP300Mut NA-ATLL cells.

### EP300 deficient NA-ATLL cells have under-replicated DNA and persistent genomic instability that results in micronuclei formation

Next, we wanted to determine whether the defective replication-associated ssDNA gaps and the DNA damage observed in EP300Mut S-phase cells persisted upon transition of cells into G2/M. Perturbed DNA replication has been associated with incompletely replicated DNA, visualized as ultra-fine DNA bridges (UFB) during mitosis. UFB’s that originate from CFS loci can be identified by twin foci formed by the Fanconi anemia complementation group D2/I (FANCD2/FANCI), proteins at the termini of the UFB on each chromatid^54^. To determine the presence of under-replicated DNA in NA-ATLL cells, we measure the persistence of FANCD2 twin foci in mitotic cells. This analysis revealed that EP300Mut NA-ATLL cells have a striking spontaneous increase in FANCD2 twin foci in mitotic cells, indicating incompletely replicated DNA arising from CFS loci as the primary source of post-replicative distress in the absence of FANCD2 (**Figure 5A**). The condensation of incompletely replicated regions, triggers the completion of DNA replication during G2/M by the process of Mitotic DNA Synthesis (MiDAS), predominantly at CFS^55,56^. MiDAS requires the coordinated activity of the nucleases, SLX4, RAD52 and the non-catalytic sub-unit of Polymerase delta, POLD3^57-59^. Analysis of MiDAS in NA-ATLL cells has revealed an increase in localization of EdU synthesis signal sandwiched between FANCD2 twin-foci in all NA-ATLL cells, irrespective of p300 expression (**Figure 5B**). However, EP300Mut cells showed a significant increase in EdU incorporation as compared to EP300WT cells. These results suggest that EP300 deficiency is associated with an increased reliance on replication completion in G2/M. Since under replication DNA and lagging chromosomes leads to micronuclei formation in the subsequent cell cycle, we next measured the percentage of cells with micronuclei positively stained for pRPA (marker of damage at collapsed forks) and FANCD2, a marker of under-replication DNA in G2/M. As expected, since NA-ATLL cells with mutated EP300 progress much slowly through cell cycle, they present with fewer numbers of micronuclei as compared to EP300WT NA-ATLL cells, however, the micronuclei observed in NA-ATLL EP300Mut cells have a prominent and significant increase in both pRPA staining and FANCD2 foci (**Figure 5C-D**), clearly indicating replication defects as the source of micronuclei formation in the absence of p300.

**Figure 5:**
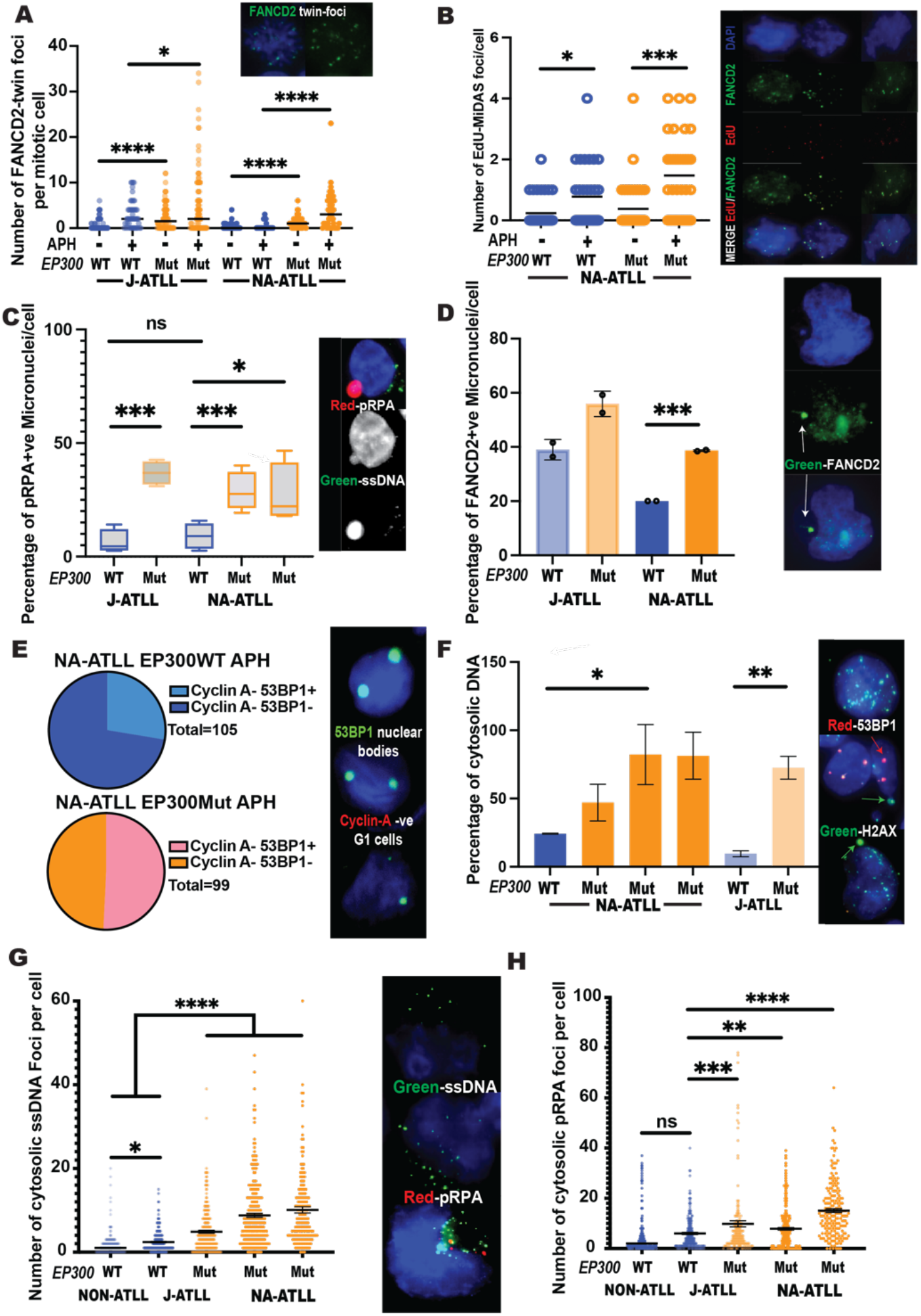
EP300 mutated cells have under-replicated DNA that results in inherited DNA lesions in G1 and manifest cytoplasmic DNA. A. Analysis of number of FANCD2 twin-foci (green) per mitotic cell nucleus (DAPI, blue) in EP300Wt and EP300Mut cells exposed to Aphidicolin (APH), representative images are shown on the top, n=100; B. Analysis of number of EdU foci (red), sandwiched between FANCD2 twin-foci, per mitotic cell nucleus (DAPI, blue) in EP300Wt and EP300Mut cells exposed to Aphidicolin (APH), representative images are shown on the right, n=100; C. Analysis of the percentage of pRPA (red) positive micronuclei in EP300Wt and EP300Mut cells, in the absence of any inhibitor treatments. Representative images are on the right, n=100 micronuclei; D. Percentage of FANCD2 (green) positive micronuclei in EP300Wt and EP300Mut cells, in the absence of any inhibitor treatments. Representative images are on the right, n=100 micronuclei; E. Analysis of number of 53BP1 nuclear body positive G1 cells (cyclin A negative) in EP300Wt and EP300Mut cells exposed to Aphidicolin (APH), representative images are shown on the right, n∽150; F. Percentage of cytoplasmic chromosome fragments (CCF) (+veGreen; -veRed) in EP300Wt and EP300Mut cells, representative images are shown on the right, n=150; G. Analysis of number of ssDNA foci (green) in the cytoplasm surrounding each cell nucleus in EP300Wt and EP300Mut cells. Representative images are on the right, n=250; H. Analysis of number of pRPA Ser4/8 foci (red) in the cytoplasm surrounding each cell nucleus in EP300Wt and EP300Mut cells. Representative images are on the right, n=250. The p-values are indicated as follows: * <0.03, ** <0.0021, *** <0.0002, **** <0.0001. Scale bar 10 μm.

### EP300 mutated cells inherit DNA lesions in G1 and manifest cytoplasmic DNA

Given the elevated genomic instability in EP300Mut mitotic cells characterized by damage containing micronuclei, we assessed mitotic mis-segregation defects into the subsequent cell cycle. Persistent unresolved DNA damage in newly formed daughter cells are localized to 53BP1 nuclear bodies in G1^60,61^. To evaluate whether unresolved DNA damage in NA-ATLL cells is transmitted as inherited lesions, we assessed 53BP1 nuclear bodies by immunofluorescence staining in cyclin A negative G1 cells. This analysis revealed a significant increase in 53BP1 nuclear bodies in G1, in the absence of EP300 (**Figure 5E**). Next, to evaluate whether these outcomes result in endogenous cytosolic DNA accumulation in EP300Mut cells, we assessed the presence of cytosolic DNA called cytoplasmic chromosome fragments (CCF). Endogenous cytosolic DNA are shown to occur in many cancers^62^. Cytosolic DNA of this nature are identified by positive staining for γH2AX and negative staining for 53BP1. The results from this analysis revealed a highly elevated presence of γH2AX (green) positive CCF in NA-ATLL EP300Mut cells (**Figure 5F**). These results were confirmed by the accumulation of mis-localized ssDNA foci and DNA stained positive for pRPA4/8 foci in the cytosol preferentially in EP300Mut cells. Taken together, these results suggest that unresolved DNA damage arising from replicative defects in EP300 deficient cells, lead to highly toxic cytosolic DNA.

### EP300 mutated cancers closely resemble BRCA-deficient tumors

In normal cells, replication fork stalling in response to replication inhibition (depletion of nucleotide pools, DNA polymerase inhibition, oncogene activation), results in fork reversal at nascently synthesized DNA followed by RAD51 loading and the efficient restart of stalled forks in the presence of downstream effectors such as BRCA1/BRCA2/ PALB2, thus maintaining genome stability (**Figure 6**, top). However, in the absence of the p300 acetyl transferase, elevated endogenous replication stress results in dormant origin firing at common fragile sites, and increased genome-wide replication pausing. While fork regression followed by RAD51 leading occurs in these cells, defective downstream effectors, such as BRCA2, lead to excessive nucleolytic degradation at regressed forks resulting in increased ssDNA at collapsed forks. This in turn triggers POLD3-mediated replication restart during the S-phase, collectively mimicking phenotypes observed in BRCA-deficient cancers. Upon transitioning to G2/M, p300 deficient cells rely on POLD3 dependent MiDAS to overcome under-replicated DNA which is largely insufficient to repair the excessive persistent damage, resulting in miss-segregation of DNA during mitosis to daughter cells in G1 and the increased accumulation of toxic ssDNA in the cytosol (**Figure 6**, bottom). Taken together, the results from this study indicate that the p300 protein plays an important previously undiscovered role in replication fork protection, that is essential to maintaining genome stability.

**Figure 6:**
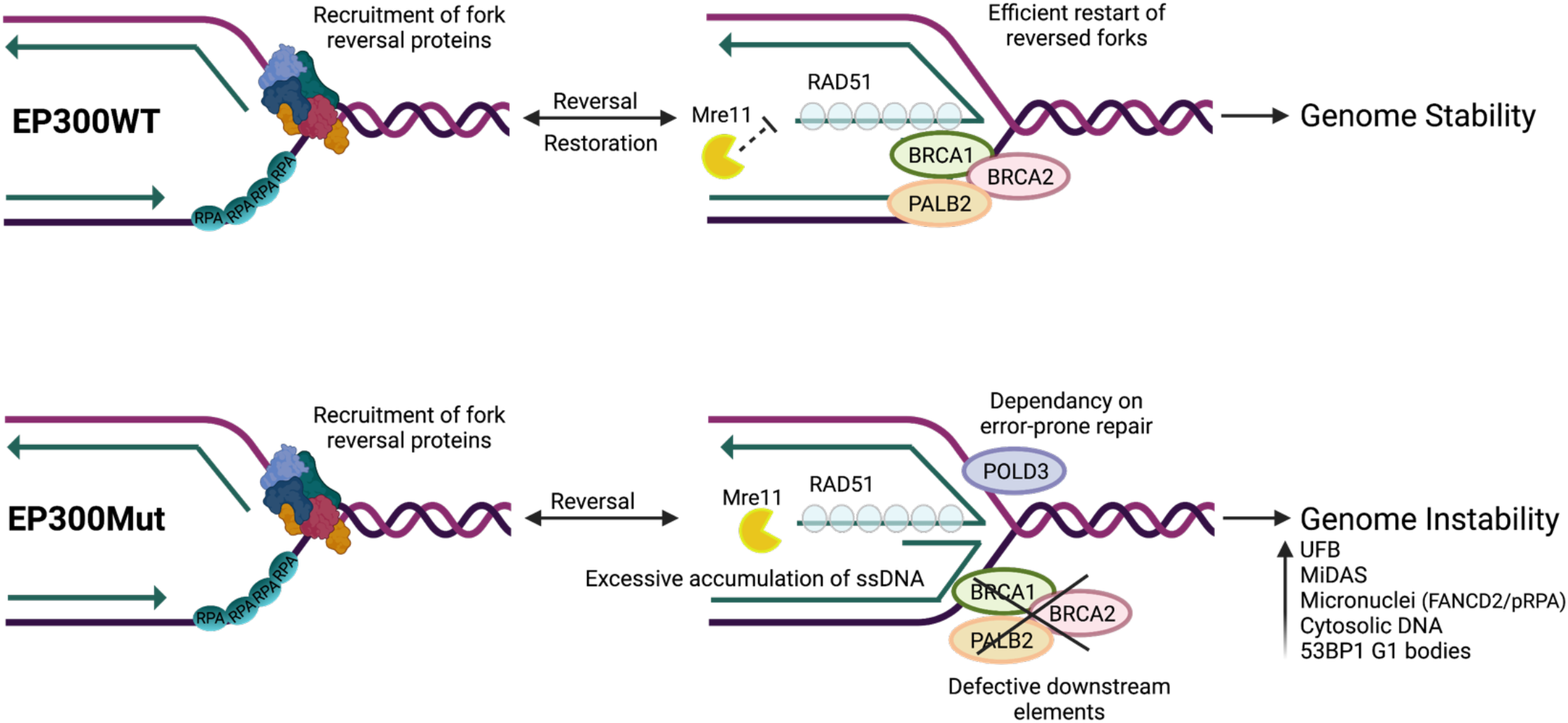
Schematic Model of Replication Fork Instability in the Absence of EP300. **Top**, In EP300WT cells, replication fork stalling in response to replication inhibition (depletion of nucleotide pools, DNA polymerase inhibition, oncogene activation), results in fork reversal at nascently synthesized DNA followed by RAD51 loading and the efficient restart of stalled forks in the presence of downstream effectors such as BRCA1/BRCA2/ PALB2, thus maintaining genome stability. **Bottom**, However, in EP300Mut cells, elevated endogenous replication stress results in dormant origin firing at common fragile sites, and increased genome-wide replication pausing. While fork regression followed by RAD51 leading occurs in these cells, defective downstream effectors, such as BRCA2, lead to excessive nucleolytic degradation at regressed forks resulting in increased ssDNA at collapsed forks. This in turn triggers POLD3-mediated replication restart during the S-phase, collectively mimicking phenotypes observed in BRCA-deficient cancers. Upon transitioning to G2/M, p300 deficient cells rely on POLD3 dependent MiDAS to overcome under-replicated DNA which is largely insufficient to repair the excessive persistent damage, resulting in miss-segregation of DNA during mitosis to daughter cells in G1 and the increased accumulation of toxic ssDNA in the cytosol.

## Methods

### Cell Culture

Human cell line GM03798 (wild type), Epstein–Barr virus-transformed lymphoblasts were obtained from Coriell Cell Repositories and were grown in RPMI 1640 (Gibco) supplemented with 10% FBS and penicillin/streptomycin (Gibco). Japanese ATL43+ and ATL43-cells were cultivated in RPMI 1640 medium supplemented with 10% fetal bovine serum (FBS), 1% penicillin/streptomycin, and with or without interleukin 2 (IL-2), respectively. North American ATLL cells were cultivated in IMDM medium supplemented with 20% human serum, 1% penicillin/streptomycin, and 100 unit/ml IL-2. Cells were treated with the different concentrations of the following drugs: Hydroxyurea (H8627-1G; Sigma); Aphidicolin (A0781, Sigma); Camptothecin (ab120115; Abcam). Cell lines were regularly tested for mycoplasma contamination.

### DNA fiber analysis

DNA fibers were stretched and prepared using a modification of a procedure described previously^63,64^. Briefly, cells were pulse labeled with 30 μM IdU/CIdU for 20 minutes each. Labelled cells were resuspended in cold PBS at 1 × 106 cells/ml. Two μl of cell suspension was spotted onto a clean glass slide and to lyse it, 10 μl of spreading buffer (0.5% SDS, 200 mM Tris-HCl (pH 7.4), and 50 mM EDTA) was added. The cells were incubated for 6 minutes, and the slides were tilted to spread the DNA. Slides were either fixed in methanol and acetic acid (3:1) for 2 min, followed by denaturation with 2.5 M HCl for 30 min at room temperature. Alternatively, the DNA was denatured with sodium hydroxide in ethanol and then fixed with glutaraldehyde. The slides were blocked with 1% BSA for at least 20 min. The slides were incubated with the antibodies as described above for SMARD. The coverslips were mounted with ProLong gold antifade reagent (Invitrogen) after a final PBS/CA630 rinse. Fluorescence microscopy was carried out using a Zeiss microscope to monitor the IdU/CIdU nucleoside incorporation.

### Immunofluorescence

Suspension cells adhered to Poly-L-lysine slides were fixed and permeabilized simultaneously in PTEMF buffer (20 mM PIPES pH 6.8, 10 mM EGTA, 0.2% Triton X 100, 1 mM MgCl2 and 4% formaldehyde). Slides were then incubated with primary antibody for Phospho-Histone H2A.X (Ser139) (Cell Signaling Technology) overnight at 4°C, and then with an Alexa Fluor 488 secondary antibody (Cell Signaling Technology) for 60 min at RT. After the secondary antibody incubation, slides were washed three times with 1X PBS and Click-It chemistry was utilized to detect EdU incorporation according to the manufacturer’s instructions (Click-IT EdU; Alexa fluor 594 Imaging Kits, Life Technologies). Slides were then mounted with Vecta Shield with DAPI (Vector Laboratories). Images were captured using a Zeiss fluorescence microscope.

### Western Blotting

Cells were harvested after treatment by centrifugation followed by PBS wash. Cells were lysed with 2X Laemmli buffer and lysates were denatured at 100ºC for 15 minutes. Proteins were separated using NuPAGE 4-14% Bis-Tris Mini Gels (Thermo Scientific) or 3-8% Tris-Acetate mini gels (Thermo Scientific) and transferred to nitrocellulose membrane. Membranes were blocked with AdvanBlock (Advansta) blocking buffer for 1 hour at room temperature and then incubated with primary antibodies overnight at 4ºC. Membranes were washed 3 times for 15 minutes and then incubated with horseradish peroxidase (HRP)-linked secondary antibodies for 1 hour at room temperature. Proteins were then detected by chemiluminescence. Time Course Experiment: Cells were grown in corresponding media and treated with 2mM HU for 4 hours. Cells were released from HU into fresh media and collected at various time points after release: 0, 4, 8, 12,24, 28, 32, and 48 hours after release. Cells were then harvested and lysed for immunoblotting. Antibodies for western blot included anti-Vinculin (Sigma Aldrich – V9131), anti-yH2AX (Cell Signaling – 9718T), anti-pCHK1 (s317) (Cell signaling –123402), anti-pRPA (ser4, ser8) (Bethyl Lab A300-245A-M), anti-CHK1 (Cell Signaling - 2360), anti-RPA (Bethyl – A300-244A), anti-H3 (Cell Signaling – 12648T), anti-H3K27ac (Cell Signaling 8173T), anti-EP300 (ABCAM-ab10485), anti-BRCA1 (EDM Millipore – 07434), anti-BRCA2 (EDM Millipore – OP95), anti-rad51 (Abcam – ab63801).

### Single Molecule Analysis of Replicated DNA (SMARD)

SMARD analysis was carried out using a previously described procedure^37,65,66^. Briefly, exponentially growing cells were cultured in media containing 30 μM 5-iodo-2′-deoxyuridine (IdU) at 37°C for 4h (Sigma-Aldrich, St. Louis, MO). After 4h, the cells were centrifuged at 800rpm for 5 min and the media containing IdU was removed. The cells were then cultured in fresh RPMI medium containing 30 μM 5-chloro-2′-deoxyuridine (CIdU) (Sigma-Aldrich, St. Louis, MO) and incubated for an additional 4h. After 4h, the cells were collected by centrifugation, and were resuspended at 3 × 107 cells per ml in PBS. The cells were resuspended in an equal volume of molten 1% InCert agarose (Lonza Rockland, Inc., Rockland, ME) in PBS. DNA gel plugs were made by pipetting the cell-agarose mixture into a chilled plastic mold with 0.5- x 0.2-cm wells with a depth of 0.9 cm. The gel plugs were allowed to solidify on ice for 30 min. The cells in the plugs were lysed in buffer containing 1% n-lauroylsarcosine (Sigma-Aldrich), 0.5 M EDTA, and 20 mg/ml proteinase K. The gel plugs were incubated at 50°C for 3 days and treated with fresh proteinase K at 20 mg/ml concentration (Roche Diagnostics), every 24 h. The Proteinase K digested plugs were then rinsed in Tris-EDTA (TE) and subjected to phenylmethanesulfonyl fluoride (PMSF) (Sigma-Aldrich) treatment. To prepare the cells for restriction enzyme digestion, the plugs were washed with 10 mM MgCl2 and 10 mM Tris-HCl (pH 8.0) and the genomic DNA in the gel plugs was digested with 70 units of PmeI or SbfI (New England BioLabs Inc.) at 37°C overnight. The digested gel plugs were rinsed with TE and cast into a 0.7% SeaPlaque GTG agarose gel (Lonza Rockland, Inc.) for size separation of DNA by pulse field gel electrophoresis. Gel slices from the appropriate positions in the pulsed-field electrophoresis gel were melted at 72°C for 20 min. The melted agarose was digested with GELase enzyme (Epicentre Biotechnologies 1 unit per 50 μl of agarose suspension) by incubating the GELase-DNA-agarose mixture at 45°C for 4 h. The resulting DNA was pipetted along one side of a coverslip that had been placed on top of a 3-aminopropyltriethoxysilane (Sigma-Aldrich)-coated glass slide and allowed to enter by capillary action. The DNA was denatured with sodium hydroxide in ethanol and fixed with glutaraldehyde.

The slides containing the DNA were hybridized overnight with biotinylated probes (represented as blue bars on the CFS locus maps). The next day, the slides were rinsed in 2 × SSC (1× SSC is 0.15 M NaCl plus 0.015 M sodium citrate) 1% SDS and washed in 40% formamide solution containing 2 × SSC at 45°C for 5 min and rinsed in 2 × SSC-0.1% IGEPAL CA-630. Following several detergent rinses (4 times in 4× SSC-0.1% IGEPAL CA-630), the slides were blocked with 1% BSA for at least 20 min and treated with Avidin Alexa Fluor 350 (Invitrogen Molecular Probes) for 20 minutes. The slides were rinsed with PBS containing 0.03% IGEPAL CA-630, treated with biotinylated anti-avidin D (Vector Laboratories) for 20 min, and rinsed again. The slides were then treated with Avidin Alexa Fluor 350 for 20 min and rinsed again, as in the previous step. The slides were incubated with the IdU antibody, a mouse anti-bromodeoxyuridine (Becton Dickinson Immunocytometry Systems), the antibody specific for CldU, a monoclonal rat anti-bromodeoxyuridine (anti-BrdU) (Abcam) and biotinylated anti-avidin D for 1 h. This was followed by incubation with Avidin Alexa Fluor 350 and secondary antibodies, Alexa Fluor 568 goat anti-mouse IgG (H+L) (Invitrogen Molecular Probes), and Alexa Fluor 488 goat anti-rat IgG (H+L) (Invitrogen Molecular Probes) for 1 hour. The coverslips were mounted with ProLong gold antifade reagent (Invitrogen) after a final PBS/CA630 rinse. Fluorescence microscopy was carried out using a Zeiss fluorescence microscope to monitor the IdU/CIdU nucleoside.

### Comet Assay

Cells were collected by centrifugation at 1000rpm for 5 minutes. Samples were then washed once with 1X PBS and centrifuged again at 1000rpm for 3 minutes. Samples were then resuspended in 1mL of 1X PBS at 1×10^6^ cells/mL. On slides coated with 1% normal melting point agarose, a mixture of 1:7 cell suspension to 1% low melting point agarose was added to each slide and covered with a coverslip. Slides were placed in 4°C for thirty minutes. Cover slips were gently removed, and cells were placed in lysis buffer (5M NaCl, 0.5M EDTA, 10mM Tris-HCl ph10, 1% Triton) for 90 minutes at 4°C. Lysis buffer was removed and denaturation buffer (300mM NaOH, 1mM EDTA) was added for 30 minutes at 4°C. Denaturation buffer was removed, and slides were placed into an electrophoresis unit containing 1X TBE buffer. Electrophoresis was run at 3V/cm for 30 minutes at room temperature. Slides were removed from the electrophoresis unit and put into 2 washes with distilled water for 5 minutes each. Slides were then dehydrated using washes with ice cold 70% ethanol 3 times for 5 minutes each.

Slides were left to dry at room temperature and stained with 1X Vista Green dye. Images were acquired using 20X magnification on Zeiss Axio microscope. Data was analyzed using OpenComet plugin via ImageJ.

### Non-Denaturing ssDNA gap detection by immunofluorescence

Cells were grown in the presence of 50uM IdU for 48 hours before treatment with 2mM hydroxyurea. Cells were then collected by centrifugation and washed with PBS (Phosphate Buffered Saline). Cells were seeded in glass slides by cytospin for 6 minutes at 1100 RPM. Cells were fixed and permeabilized with 0.5% Triton X-100 and 4% paraformaldehyde in PBS at room temperature for 15 minutes. Fixed cells were then blocked with 3% BSA for 1 hour at room temperature. Cells were incubated with primary antibodies against pRPA at 4ºC overnight. Cells were then washed 3 times with PBS at room temperature followed by incubation with antibodies against IdU for 1 hour at room temperature. Cells were washed 3 times with PBS followed by secondary antibodies for 1 hour at room temperature. Cells were then mounted with Prolong with DAPI.

### Mitotic DNA Synthesis (MiDAS)

Harvesting mitotic cells: To augment mitotic cells, asynchronous cells were treated with CDK1 inhibitor RO-3306 (Merck) at a final concentration 7 μM for 7:30 h, or in the last 7:30h of APH treatment as and when needed. The cells were then washed with pre-warmed Edu-containing medium (37 °C) for three times within 5 minutes before being released into pre-warmed fresh medium with Edu at a final concentration of 20 μM for 45mins-1 h. This allows the cells to advance into mitosis. The mitotic population is then analyzed. Next, the cells are collected and centrifuged at RT at 350g/rcf for 5 minutes to form a pellet. They are then washed in pre-warmed fresh medium two times before resuspending them into DPBS (1 million cells/ml of DPBS). Further, cells are seeded on Corning-single frosted microslides using Cytospin 4 (Thermoscientific) for 6 minutes at 1100rpm at medium acceleration. Once cells are seeded, they are fixed and permeabilized for 15 minutes with 0.5% Triton in DPBS containing 4% formaldehyde. The slides are then washed twice with DPBS- to be used for EdU labelling and IF, or they can be stored at 4 °C for maximum two weeks.

EdU detection and IF staining for MIDAS: The cells were blocked using blocking buffer (3% BSA in 1X PBS) for 2 hours at RT. To detect EdU Click-IT chemistry was used as per the manufacturer’s instructions (Click-iT™ EdU Cell Proliferation Kit for Imaging, Alexa Fluor™ 594 dye). EdU detection was coupled with IF staining. The cells were incubated with primary antibody diluted in the blocking buffer at 4 °C overnight; followed by three washes 15 mins each with DPBS. Next, the cells were incubated with secondary antibodies diluted in the blocking buffer for 1 hour at RT, followed by three washes, 15 mins each with DPBS. Slides were then mounted with Prolong gold containing DAPI (Invitrogen Prolong Gold anti fade reagent with DAPI, P36935). The slides were allowed to dry overnight prior to IF imaging. Images were captured using-63mm oil immersion lens, Ziess Axio microscope.

Primary and Secondary antibodies for MiDAS experiments: Primary antibodies used were FANCD2 (1:1000, NB100-182, Novus), MUS81 (1:250, ab14387, abcam), RAD52 (1:250, sc-365341, Santa Cruz), 53BP1 (1:5000, S1778, Cell Signaling or NB100-304,Novus), Cyclin-A(1:500, sc-271682, Santa Cruz), POLD3 (1:1000, H00010714-M01, abnova). Secondary antibodies used were Anti-rabbit IgG Fab2 Alexa Fluor ® 488 (1:20000, 4412S, Molecular probes), were Anti-rabbit IgG Fab2 Alexa Fluor ®594 (1:20000, 8889S, Molecular probes), Anti-mouse IgG Fab2 Alexa Fluor ® 488 (1:20000, 4408S, Molecular probes), Anti-mouse IgG Fab2 Alexa Fluor ® 594 (1:20000, 8890S, Molecular probes).

### Analysis

#### Under-replicated DNA at CFS

Mitotic cells were labelled for EdU and IF staining was performed. FANCD2 twin foci were used as a marker of CFS location. Each mitotic cell was analyzed for EdU labelled FANCD2 twin foci. Each FANCD2 twin foci, showed 1-2 EdU foci suggesting DNA replication at under-replicated regions called CFS.

#### Inherited Nuclear lesions

Cells arrested at G2 using RO-3306 (Merck) at 7μM final concentration were released and allowed to continue with cell cycle for 1 hour. After fixing, blocking and IF staining, and imaging, the cells were classified into Cyclin A positive and Cyclin A negative population. Cyclin A negative cells stained for distinct 53BP1 foci are analyzed as G1 nuclear lesions.

#### Micronuclei Analysis

Chromatin fragment staining positive for DAPI, proximal to parent nucleus was analyzed as micronucleus. These micronuclei were stained for proteins that mark damaged regions as well as for sensing cytosolic DNA. The DAPI stained micronuclei that are stained for H2AX, 53BP1 and pRPA were considered indications of damage response. They were further classified into Micronuclei and CCFs (Cytoplasmic Chromatin Fragments). CCFs were specifically the fragments that stained positive for H2AX but were negative for 53BP1.

### Quantification and Statistical Analysis

All statistical calculations were performed using GraphPad Prism v.9.0 A two-sided Student’s t-test was used to calculated p-values in GraphPad Prism v.9.0. All the experiments were generally conducted with at least three biological replicates. Differences were considered significant at p-values: * <0.03, ** <0.0021, *** <0.0002, **** <0.0001.

## Author Contributions

The project was conceived by A.M. Experiments most of the experiments were conducted by A.B.G. The MiDAS experiments were conducted by M.N and the Comet assay was conducted by J.G. Data analysis was carried out by A.B.G, M.N, J.G, R.Z, V.K, A.J, A.P, and I.S. The data was analyzed and discussed by A.M, A.B.G, J.G, J.C, K.K, A.D, and B.H.Y. The JQAD1 compound was provided by J.Q. The manuscript was written by A.M.

## Acknowledgements

The work was supported by an American Cancer Society pilot award to A.M and R00HL136870 to A.M.

## Author Information

The authors have NO commercial affiliations or conflicts of interest.

