## Supplemental Figures for "Acetyl transferase EP300 deficiency leads to chronic replication stress mediated by defective fork protection at stalled replication forks"

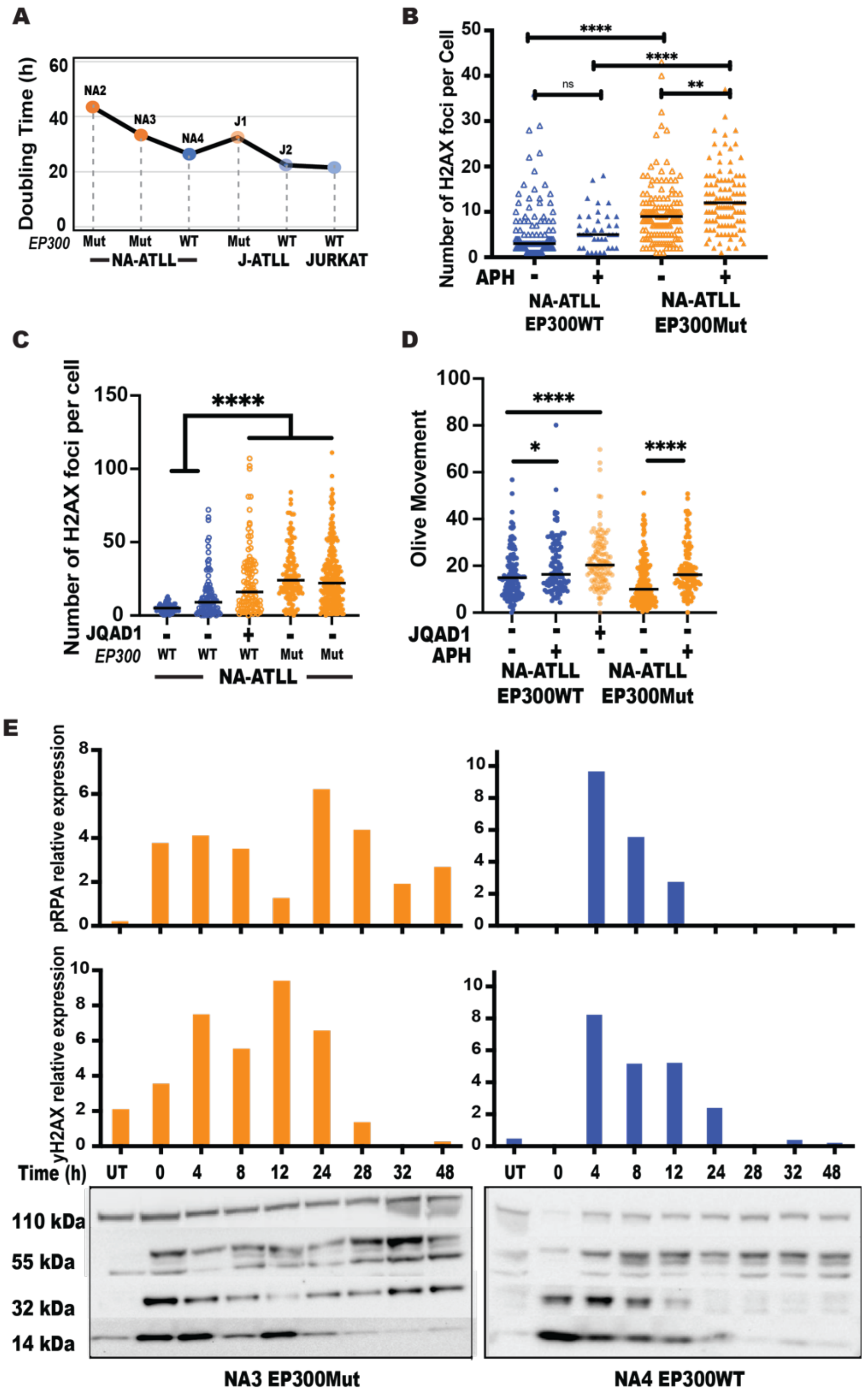


**Supplemental Figure 1: *EP300*-mutated cells have aberrant cell cycle dynamics, spontaneous DNA damage and persistent replicative checkpoint activation.** A. Calculation of cell doubling times using the open source cell doubling time calculator (https://www.omnicalculator.com/biology/cell-doubling-time). Cell doubling was calculated every 48 hours from three independent cultures of cells; B-C. Analysis of the number of pH2AX foci per cell nuclei in EP300WT cells treated with APH and(or) JQAD1, n=150; D. Measurement of DNA single strand breaks by alkaline Comet assay in EP300Mut and EP300WT cells treated with APH in the presence or absence of JQAD1. Comet tail lengths were measured using the OpenComet plugin as part of the ImageJ software, n=100. The p-values are indicated as follows: * <0.03, ** <0.0021, *** <0.0002, **** <0.0001. Scale bar 10 μm; E. Time course experiment to measure recovery of cells after release into drug free media over the course of 48hours. Cells were collected at eight time points (0, 4, 8, 12, 24, 28, 32 and 48hours) and expression levels of phospho-Chk1 Ser317, phospho-RPA Ser4/8 and phospho-Histone H2AX Ser139 were measured by western blotting. Expression levels of Vinculin was used as a loading control.

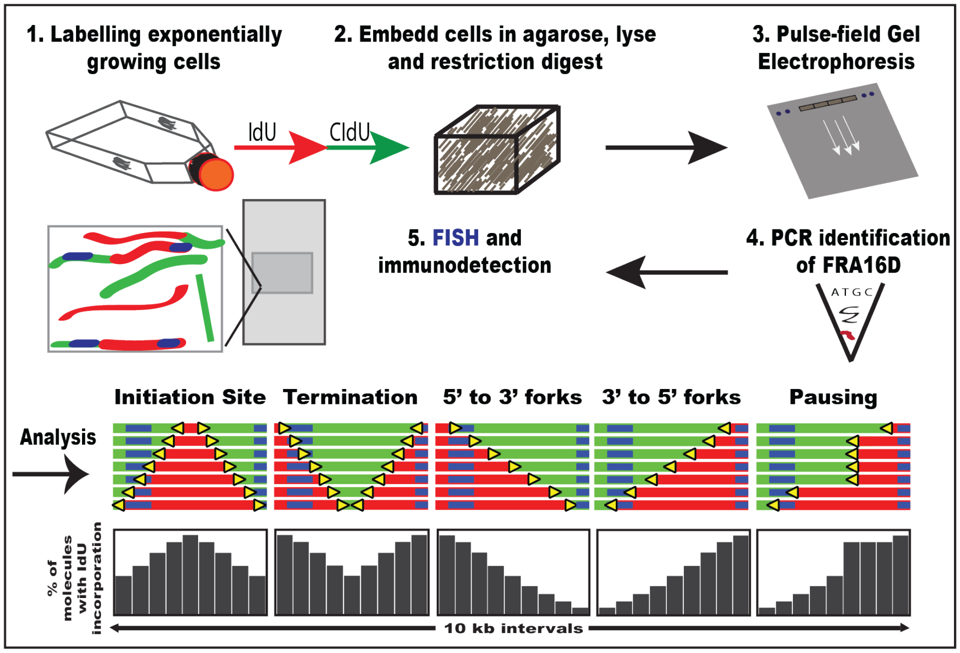


**Supplemental Figure 2: Schematic of single molecule analysis of replicated DNA (SMARD), related to Figure 2.** (A) Schematic representation of the various stages of SMARD. Cells are pulsed with nucleoside analogs (IdU-green; CIdU-red) and embedded in agarose plugs. The cells are first lysed; proteins are digested by proteinase K and then subjected to restriction digestion. The restriction digested DNA is resolved by pulse field gel electrophoresis. The slice containing the FRA16D locus is identified by PCR analysis. The agarose from the identified slice is melted and the DNA is stretched onto silanized glass slides. Biotinylated FISH probes are used for identification of fragment and immunostaining is utilized to visualize the IdU tract in red, the CIdU tract in green and the FISH probes in blue. The resulting molecules are arranged to yield recognizable replication patterns (from the left): initiating molecules, terminating molecules, replication forks travelling in the 3' to 5' and 5' to 3' direction which are easily interpreted by the IdU incorporation histograms.

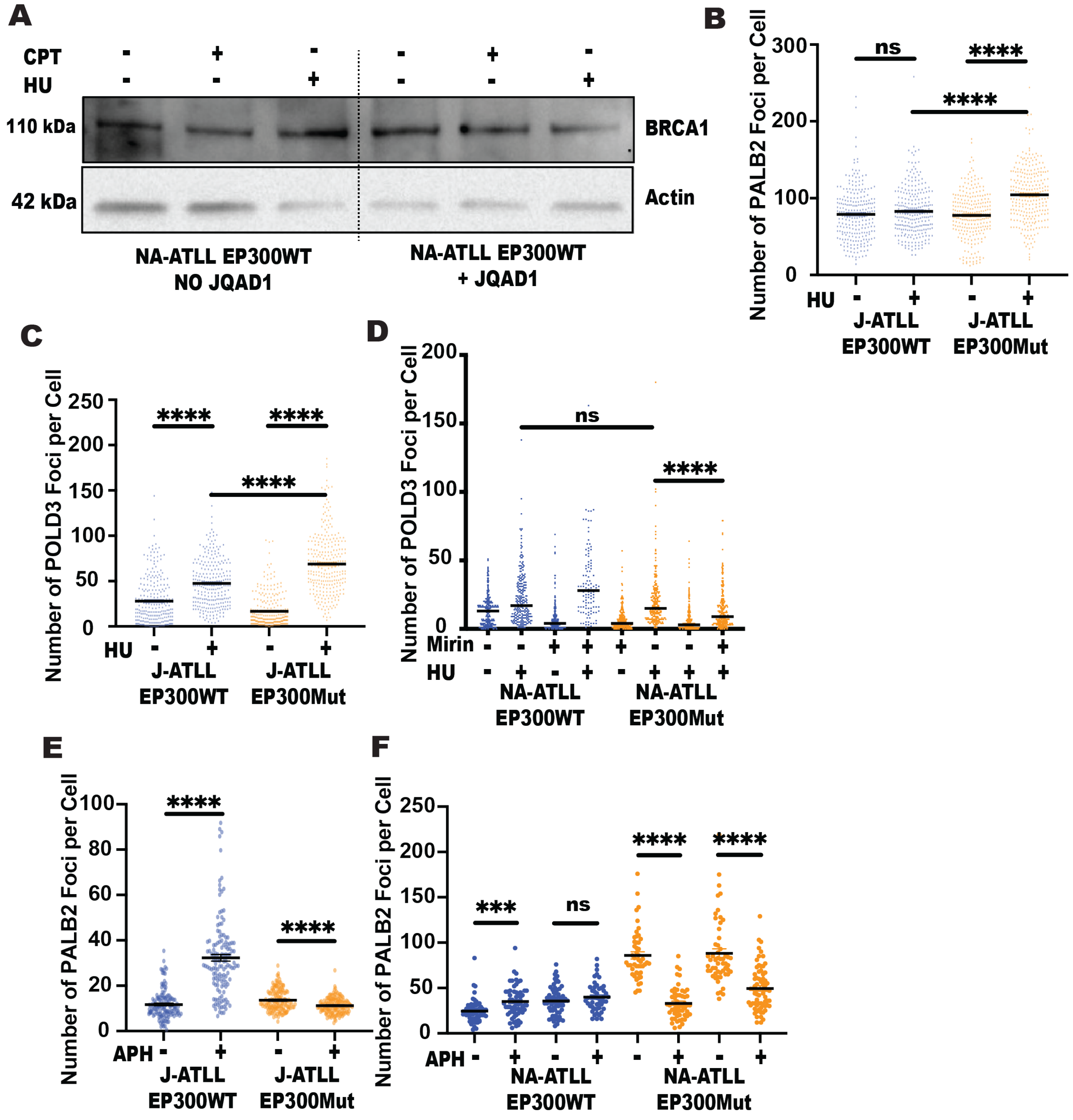


**Supplemental Figure 3: EP300 deficient cells have a prominent defect in downstream fork restart machinery**. A. Expression levels of BRCA1 protein from WCE in EP300WT NA-ATLL cells treated with HU or Camptothecin, in the presence or absence of JQAD1, by western blotting. Expression levels of Actin was used as a loading control; B. Analysis of number of PALB2 foci per cell nucleus in EP300Wt and EP300Mut NA-ATLL cells exposed to HU, n=250; C-D. Analysis of number of POLD3 foci per cell nucleus in EP300Wt and EP300Mut J/NA-ATLL cells exposed to HU, in the presence or absence of Mirin, n=250; E-F. Analysis of number of PALB2 foci per cell nucleus in EP300Wt and EP300Mut J/NA-ATLL cells exposed to APH, n=150. The p-values are indicated as follows: * <0.03, ** <0.0021, *** <0.0002, **** <0.0001. Scale bar 10 μm.
